# The role of L-arabinose metabolism for *Escherichia coli* O157:H7 in edible plants

**DOI:** 10.1101/2021.04.08.437851

**Authors:** Louise Crozier, Jacqueline Marshall, Ashleigh Holmes, Kathryn Wright, Yannick Rossez, Bernhard Merget, Sonia Humphries, Ian Toth, Robert Wilson Jackson, Nicola Jean Holden

## Abstract

Arabinose is a major plant aldopentose in the form of arabinans complexed in cell wall polysaccharides or glycoproteins (AGP), but comparatively rare as a monosaccharide. L-arabinose is an important bacterial metabolite, accessed by pectolytic microorganisms such as *Pectobacterium atrosepticum* via pectin and hemicellulose degrading enzymes. However, not all plant-associated microbes encode cell wall degrading enzymes, yet can metabolise L-arabinose, raising questions about their use of and access to the glycan in plants. Therefore, we examined L-arabinose metabolism in the food-borne pathogen *Escherichia coli* O157:H7 (isolate Sakai) during its colonisation of plants. L-arabinose metabolism (*araBA*) and transport (*araF*) genes were activated at 18 °C *in vitro* by L-arabinose and expressed over prolonged periods *in planta*. Although deletion of *araBAD* did not impact the colonisation ability of *E. coli* O157:H7 (Sakai) on plants, *araA* was induced on exposure to spinach cell wall polysaccharides. Furthermore, debranched and arabinan oligosaccharides induced *ara* metabolism gene expression *in vitro*, and stimulated modest proliferation, while immobilised pectin did not. Thus, *E. coli* O157:H7 (Sakai) can utilise pectin/AGP- derived L-arabinose as a metabolite, but differs fundamentally in *ara* gene organisation, transport and regulation from the related pectinolytic species *P. atrosepticum*, reflective of distinct plant- associated lifestyles.

## Introduction

Arabinose is an abundant aldopentose in plant material that is not found in animals. It is present almost entirely as polysaccharides (arabinans) and glycoproteins, as L-arabinofuranose (L-Ara*f*) complexed in the RG1 and RG2 pectin components of plant cell walls [1], as side chains in hemicellulose [2] or in arabinogalactan-proteins, AGP [3]. Free monomeric L-arabinopyranose (L- Ara*p*) is present in intracellular and apoplastic compartments but is rare in comparison to fructose and sucrose [4]. L-arabinose metabolism is widespread in microbes, reflective of a beneficial function for accessing the carbohydrate directly from plants or as a dietary fibre in animal guts. Microbial metabolism of L-arabinose has been well characterised for fundamental understanding of metabolic processes [5] and for biotechnological applications [6, 7].

L-arabinose metabolic systems comprise transport and metabolism genes and a master regulator. In *E. coli* and related bacteria, L-Ara*p* is transported into the cell by an ABC transporter system where AraF is the periplasmic component that binds L-Ara*p* with high affinity, AraH is the trans-membrane protein and AraG the ATP-binding component [8]. AraE is an H^+^ symporter with relatively low affinity (140-320 μM) for L-Ara*p* [9]. Intracellular L-arabinose enters the pentose phosphate pathway in a 3- step degradation pathway via AraA, an isomerase that converts it to L-ribulose; AraB, a ribulokinase that catalyses L-ribulose phosphorylation to L-ribulose-5-phosphate; and AraD, an epimerase that converts L-ribulose 5-phosphate to D-xylulose-5-phosphate [5]. Transport and metabolism genes are under the control of AraC, an activator that is triggered when complexed with L-Ara*p* [10]. Expression of the *ara* genes is under catabolite control and cyclic AMP forms a dimeric complex with CRP to co-regulate with AraC in the absence of glucose [11]. *araF* in *E. coli* has two promoters, under the control of σ^70^ and σ^S^, respectively [12, 13], and is repressed by a small RNA [14].

Shigatoxigenic / verocytotoxigenic *Escherichia coli* (STEC/VTEC), predominately serotype O157:H7, is a food-borne pathogen that can be transmitted through the food-chain by edible plants and utilise plants as secondary hosts [15]. STEC can mobilise metabolic pathways that are specific for different plant tissues [16, 17], including L-arabinose metabolism, which was induced on exposure to crude extracts of spinach leaf lysates and spinach cell wall polysaccharides [16]. Yet, *E. coli* do not encode cell-wall degrading enzymes (PCWDE) in contrast to the related members within the *Enterobacteriaceae*. The phytopathogen *Pectobacterium atrosepticum* encodes pectinases that are considered virulence factors in plant disease [18]. This raises fundamental microbial ecology questions as to whether STEC are able to exploit an apparently inaccessible metabolite in plant- microbe interactions without the aid of cell-wall degrading enzymes. Therefore, we tested the hypothesis that L-arabinose metabolism facilitates the ability of STEC to colonise plants, by comparing L-arabinose response of *E. coli* O157:H7 isolate Sakai *in vitro* and *in planta* to that of *P. atrosepticum*. Horticultural crop species (spinach and lettuce leaves, and broccoli microgreens) that are considered a risk of foodborne illness from STEC were used as the most relevant plant models. To understand the contribution of L-arabinose metabolism to colonisation of plants by STEC (Sakai), we took a reductionist approach to quantify gene expression to in response plant extracts *in vitro*, which was validated by a qualitative approach of gene expression *in planta*.

## Material &Methods

### Bacteria

Bacterial strains were: *E. coli* STEC isolate Sakai [19]; *E. coli* strain AAEC185A [20] used for cloning; *P. atrosepticum* isolate SCRI-1043 [21] and its derivative mutants *expI-* and *outD-* [22]. An *araBAD* knock-out mutant of STEC (Sakai) was constructed by allelic exchange, cloning the upstream and downstream flanking region with primers ECs0066No_for, ECs0066Ni_rev and ECs0066Ci-for, ECs0066Co_rev on *Pst*I and *Not*I and *Not*I and *Sal*I sites, respectively, into the exchange vector pTOF24 [23], with a tetracycline gene introduced into the *Not*I site for selection. The deletion mutation was confirmed by PCR and Sanger sequencing. An *araC* knock-out mutant of STEC (Sakai) was a generous gift for this study, made by lambda-Red recombination [24]. Bacteria were routinely grown with aeration in lysogeny broth (LB) or MOPS medium [25] at 37 or 18 °C (STEC), or 27 °C (Pba) supplemented with 0.2 % glucose or glycerol where indicated, 10 µM thiamine and MEM essential and non-essential amino acids (Sigma M5550 and M7145) termed rich defined MOPS (RD- MOPS). Antibiotics were included to maintain transformed plasmids at 50 μg/ml kanamycin (Kan), 25 μg/ml chloramphenicol (Cam), 10 μg/ml Tetracycline (Tet) or 50 μg/ml ampicillin (Amp).

### Gene expression analysis

Reporter plasmids were constructed in a pACYC-derived vector, carrying the *gfp+* gene, termed pKC026 [26], with promoter regions amplified from STEC (Sakai) or Pba (1043) genomic DNA with a proof-reading polymerase (Phusion, NEB) and inserted via the *Xba*I cloning site (Suppl. Table 1). A low copy vector *gfp+* reporter was generated, derived from pWSK29, by sub-cloning the *gfp+* gene plus the *ara* genes from previously generated plasmids into the Pst1 site (Suppl. Table 1). Transformed STEC (Sakai) or Pba (1043) were grown at 18 or 27 °C, respectively in RD MOPS glycerol with aeration to OD_600_ of 2.0 and sub-inoculated at 1:100 into 10 ml RD MOPS glycerol supplemented with 10 µM to 10 mM L-arabinose. Cell density and GFP fluorescence were measured at 2h intervals, from 150 µl aliquoted into a black 96-well plate and measured in a GloMax Multi Detection System (Promega) machine (excitation 490nm, emission 510-570nm). Dose-response was measured at 0, 2, 4, 6, 8 and 24 h in RD MOPS glycerol and 0, 1, 2, 3, 4, 5 h in RD MOPS glucose. Time points for analysis were selected at maximum GFP emission, since the levels reached a maximum and then decreased presumably as arabinose was depleted and metabolism switched to glycerol. GFP measurements in plant extracts were taken at 24 hour intervals over five days, from cultures inoculated into plant extracts or RD MOPS supplemented with L-arabinose or D-glucose and grown statically at 18 °C in a black multi-well plate. Four sample reps were measured per test, and the experiment repeated three times. GFP is expressed as relative fluorescent units after subtraction from the vector-only control (pKC026 or pWSK29) data and normalised for cell density to OD_600_ of 1.0. Arabinose concentrations were transformed (Log_10_) for graph plotting and linear regression analysis. Gene expression analysis was measured directed by quantitative reverse transcriptase PCR (qRT-PCR) as previously [16] from STEC (Sakai) grown in minimal M9 medium (1 x M9 salts, 1 mM MgSO_4_, 0.3 mM CaCl_2_) with 0.2 % glycerol for 48 h at 18 °C, and sub-inoculated 1:100 in minimal M9 supplemented with 0.2 % L-arabinose or glucose, or with 0.2 % oligo- / polysaccharides ((1-5)-α- linked pectin backbone +/- α-L-arabinofuranosidase treatment; (1-5)-α-L-arabinobiose (Ara2); (1,5)- α-L-arabinoheptose (Ara7), (Megazyme, Bray, Ireland) at 18 °C and sampled as indicated. Primers for STEC *araA* and *araD* (Suppl. Table 1) were tested for efficiency and removal of DNA confirmed from direct PCR as described previously [16]. PCR products were amplified with iTaq™ Universal SYBR Green Supermix (Bio-Rad) measured on a StepOnePlus™ machine (Applied Biosystems), data normalised to *gyrB* (ECs4634) and expressed as 2^-ΔΔCT^ for fold-change compared to a control situation.

### Plants & plant extracts

Spinach (*Spinacia oleracea*) cultivar Amazon (Sutton Seeds, UK) lettuce cv. Salinas (*Lactuca sativa*) (Tozer Seeds, UK), tomato (*Solanum lycopersicum* cv. Moneymaker) (Thompson & Morgan, UK), broccoli (*Brassica oleracea* var. *italica*) (Unwins, UK), and *Nicotiana benthamiana* (Hutton stocks) were grown individually in 9 cm^3^ pots in compost for colonisation assays in a glasshouse for three weeks in standard compost. Plants for root exudates or vermiculite for polysaccharide extracts were grown from surface sterilised seeds (2 % calcium hypochlorite solution) in pots containing autoclaved autoclaved rockwool (Progrow Ltd), as described previously [16]. In brief, exudates were collected from 24 plants per batch by three aqueous extractions of the rockwool, and for the polysaccharide extracts, whole roots were excised from the plant and briefly washed in SDW to remove as much of the vermiculite as possible. A similar extract was made from vermiculite only as a negative control. Leaf and root extracts were frozen in liquid nitrogen and ground to a fine powder, for which 10 g was used for pectin (CDTA-treatment) and hemicellulose (NaOH treatment) fractions from an alcohol insoluble residue, which required a heating step (50 °C) to separate the fractions [27]. All lysates/polysaccharide extracts were stored at -20 °C.

ELISA was used to quantify arabinans, as described previously [28]. In short, 100 μl volumes of 1:10 plant polysaccharide extract (in 0.1 M NaHCO_3_ buffer, pH 9.6) were incubated overnight at 4°C to coat 96-well microtitre plates (NUNC, Maxisorb) blocked with 3 % skimmed milk protein in TBS (1 h, RT) and washed three times with TBS (100mM Tris, 150mM NaCl, pH7.5). Primary antibodies (PlantProbes) were added at 1:20 dilution in 3 % skimmed milk protein in TBS (2 h, RT) and the plates washed three times in TBS, Secondary antibody, goat anti-rat horseradish peroxidase at 1:1000 was added (1 h, RT) and the plates washed three times in TBS, then developed with ABTS solution: 22 mg ABTS (Sigma Aldrich, St. Louis, USA) diluted in 100 ml of citrate buffer (50 mM sodium citrate, 0.05% H_2_O_2_, pH 4.0). Absorbance was measured at 405nm on a microplate reader (Varioskan, ThermoFisher Scientific, USA).

Pba (1043) was grown in RD MOPS glycerol for 24 h at 27 °C, and sub-inoculated into fresh RD MOPS supplemented with 1% (w/v) spinach alcohol-insoluble polysaccharide extract plus 0.2% or 0.4% glycerol, over 24 h at 27 °C. STEC (Sakai) grown were grown in minimal M9 at 18 °C supplemented with oligosaccharides (Ara2, Ara7) exactly as per gene expression analysis, and cfu measurements taken at 0, 16, 32 and 48 h on selective MacConkey agar plates.

### Plant colonisation assay

Three-week-old plants were used for leaf or root inoculations as per [4], inoculated with STEC (Sakai), diluted to OD_600_ of 0.02 (equivalent to 10^7^ cfu/ml) in SDW used to make bacteria suspensions. Leaves were dip-inoculated by submerging the foliar parts in 1 L bacterial suspension for 30 seconds, or roots were colonised by partially submerging plant pots in the suspension for 1h. The pots were then transferred to the growth chamber until sampling. Plants were sampled at 0, 5, 10 and 14 days post infection (dpi), aerial tissue removed aseptically from with a sterile scalpel, the compost removed by washing with SDW, and the roots transferred into 50 ml tubes, washed with PBS and the fresh weight determined. The tissue was macerated with a mortar and pestle, and 10- fold dilutions plated on MacConkey + Kan agar for selective detection of STEC (Sakai) over the endemic microbiota. Data was analysed and plotted in Microsoft Excel, GraphPad Prism or RStudio, and significant differences calculated by one-way ANOVA for each tissue (P > 0.05).

### Plant colonisation for microscopy

*N. benthamiana* leaves ∼ 6 cm length were infiltrated with STEC (Sakai) at 10^7^ cfu/ml or Pba (1043) 10^6^ cfu/ml in 0.5 x Murashige and Skoog (MS) medium (Sigma Aldrich) and maintained in an environmental cabinet with 16 h daylength and a continuous temperature of 21°C. Samples were imaged 4-11 days post infiltration for STEC (Sakai) and 1-4 days post infiltration for Pba (1043). Broccoli seed was surface sterilised and germinated on purple capillary matting (Grofelt, UK) watered with bacteria at 10^3^ cfu/ml in 0.5 x MS as described previously [29] and imaged 5-11 days post sowing. Inoculated cotyledons were harvested for antibody labelling 6 days post sowing, with methods adapted from [30] using microtubule-stabilising buffer (MTSB: 50 mM PIPES, 5 mM EGTA, 5 mM MgSO_4_, 4.5 mM KOH adjusted to pH 7.0 with 10M KOH) and blocking buffer (2 % Albumin fraction V BSA in MTSB). Individual cotyledons were cut and placed in SDW, before incubation in blocking buffer for 2 h at 37°C, followed by overnight incubation at 4°C with α-O157 (*E. coli* antisera O157 monovalent, rabbit-derived, MAST ASSURE, M12030) diluted 1:500 in blocking buffer. The samples were rinsed twice in MTSB and incubated at 37°C with Alexa Fluor^®^ 568 nm goat anti rabbit (Invitrogen A11011) diluted 1:1000 in blocking buffer followed by three five-min washes and mounting in MTSB for imaging. For LM6 labelling, cotyledons were fixed in 100 % methanol for 20 min at 37°C followed by the addition of fresh methanol at 60 °C for three min, then gradual addition of water until the methanol was diluted to 20 %. The cotyledons were then transferred to fresh SDW prior to antibody labelling as described above with LM6 (PlantProbes 1,5-arabinan, 1:50) and α-O157 (1:500) followed by secondary antibodies Alexa Fluor^®^ 488 nm goat anti rat (Invitrogen A11006, 1:100) for 1 h and Alexa Fluor^®^ 568 nm goat anti rabbit (Invitrogen A11011, 1:100) for a further 2 h.

### Confocal microscopy

*N. benthamiana* leaves were infiltrated with sterile distilled water (SDW) and secured to microscope slides with double-sided adhesive tape prior to imaging of the abaxial surface as described previously [31]. Broccoli cotyledons were mounted on slides under cover-slips, in SDW for STEC (Sakai) or dry for Pba colonised leaves (since addition of water affects distribution of Pba rendering images meaningless), and imaged via either surface. Images were collected at Nyquist resolution using a Nikon A1R confocal laser scanning microscope fitted with either an NIR Apo 40x 0.8W water dipping lens or a CFI Plan Fluor 10x 0.3 lens, and GaAsP detectors. Images represent false-coloured maximum intensity projections, unless otherwise stated, produced, linearly adjusted and re-sized using the splines option, where necessary for publication, using NIS-elements AR software. GFP, Alexa Fluor^®^ 488 nm (green) and chlorophyll (blue) were excited at 488 nm with emissions collected at 500-530 nm and 663-737 nm respectively and Alexa Fluor^®^ 568 nm (magenta) excited sequentially at 561 nm with emission at 570-620 nm.

## Results

### Arabinose metabolism genes are induced on exposure to spinach extracts

Whole transcriptomic analysis of *E. coli* O157:H7 (Sakai) from a previous study had shown induction of arabinose-associated genes on exposure to spinach extracts (Table 1, extracted from [16]). The differential expression profiles were confirmed using a different approach (qPCR) and showed that the isomerase, *araA* was induced on exposure to spinach cell wall polysaccharides but not lettuce (Supplemental Fig. 1a). In contrast, *araA* expression was reduced after 1 h following infiltration into either spinach leaves (16.82-fold +/- 4.68) or lettuce leaves (14.08-fold +/- 1.89) relative to the no-plant control condition (Supplemental Fig. 1b). Thus, STEC (Sakai) response to L-arabinose appears dynamic and is impacted by tissue type, raising the hypothesis that L-arabinose metabolism impacts STEC (Sakai) colonisation of plants.

**Table 1.**
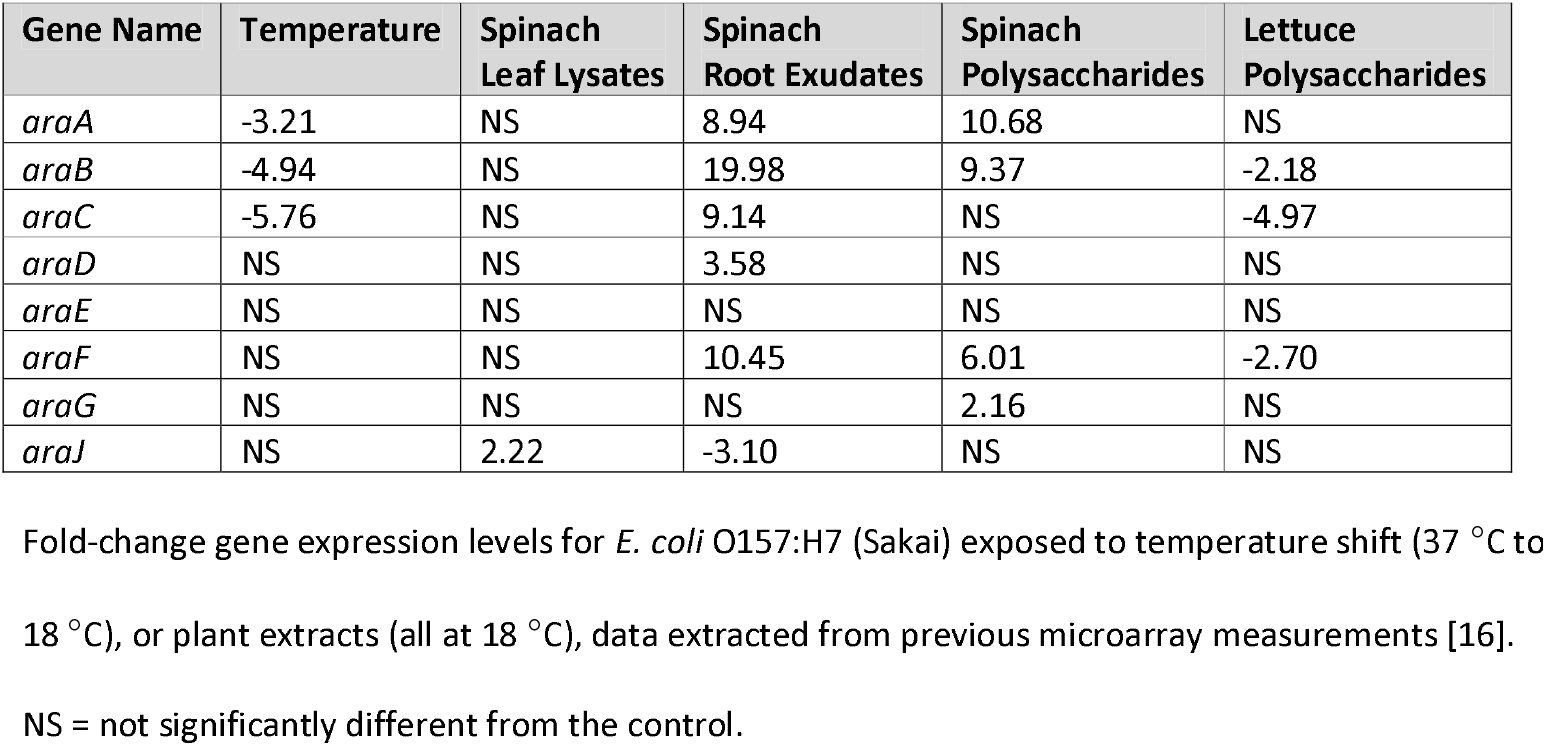
Expression of genes related to the processing, transport and utilisation of arabinose.

### Arabinose metabolism is not essential for colonisation of plants

To determine any essentiality of arabinose metabolism *in planta*, a knock-out mutant of the *araBAD* operon was made in STEC (Sakai) and tested in plant colonisation ability primarily on spinach leaves and confirmed on the original plant models, spinach roots, lettuce roots and leaves. Both the WT and the *araBAD* mutant strains showed essentially the same patterns of colonisation, on both species and in both tissue types, with a decrease from the initial starting inoculum over time, which was greatest on the foliar tissue, as expected from a high inoculation dose. There was no significant difference between the WT and *araBAD* mutant (> 95 %) and any reductions in colonisation by the *araBAD* mutant strain were marginal, e.g. on lettuce roots at day 10 (Fig. 1). Thus, the arabinose metabolism genes are not essential for STEC (Sakai) colonisation of spinach or lettuce, leaves or roots.

**Figure 1.**
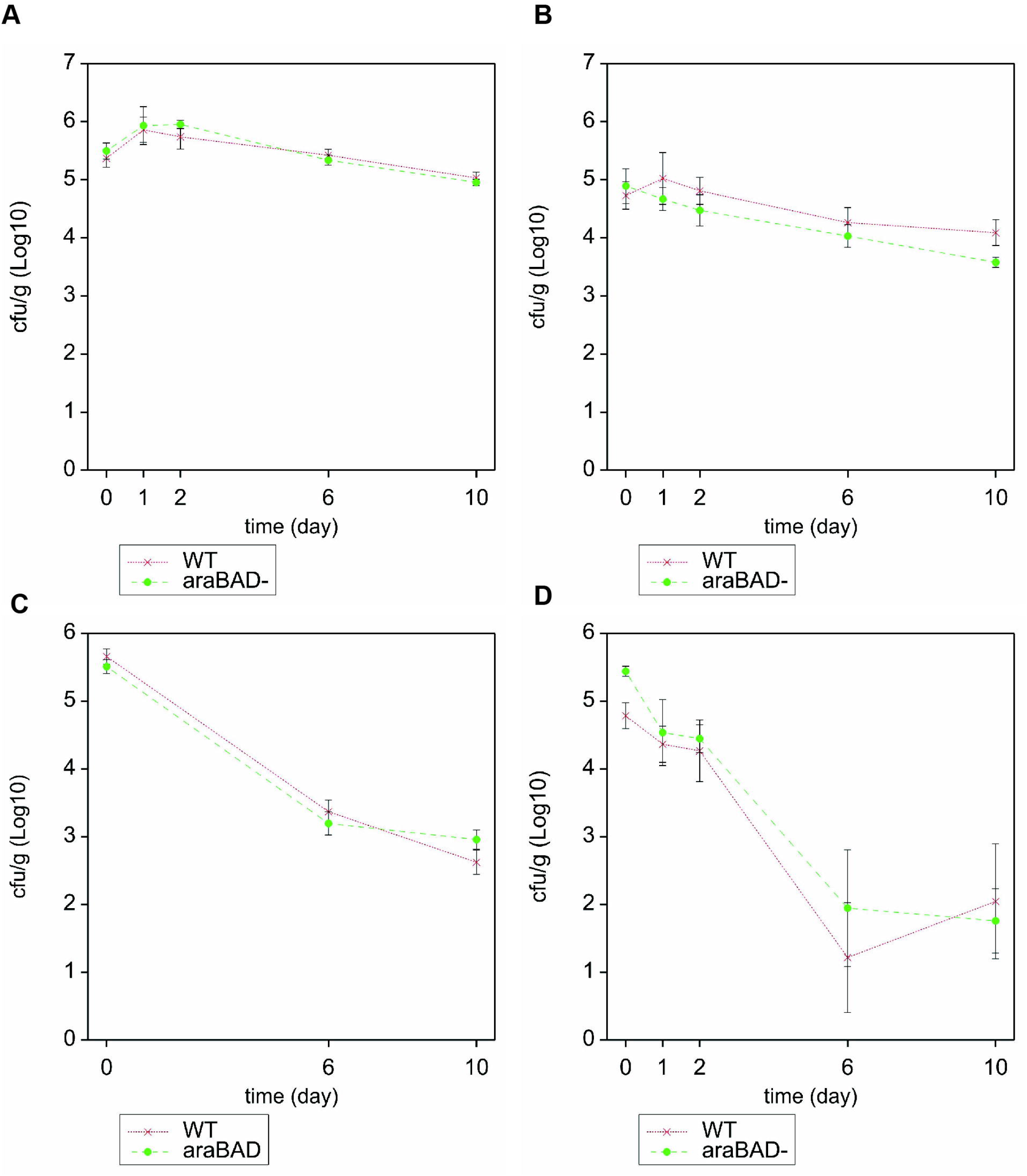
Colonisation of spinach and lettuce plants by Sakai WT and Sakai *araBAD*. Number of bacteria recovered from spinach (A, C) or lettuce (B, D) roots (A, B) or leaves (C, D). The averaged (+/- SD) are shown for STEC (Sakai) WT (red) or *araBAD* mutant (green), enumerated from four (lettuce) or five (spinach) plant samples and the experiment was repeated three times for spinach leaves. Samples were taken on day 0, 1, 2, 6 and 10 or for spinach leaves on day 0, 6, 10.

### Transcriptional activation of STEC (Sakai) ara genes in vitro

Since arabinose metabolism was not essential for STEC colonisation of plant tissue, expression of STEC (Sakai) *ara* metabolism and transport genes was examined in detail under *in vitro* conditions. Transcriptional GFP reporter plasmids were generated for metabolism (*araBA*) and high-affinity transport (*araFGH*) genes of STEC (Sakai). Responsiveness to L-Ara*p* was measured at 18°C, a temperature relevant to colonisation of plants, and to show utility of the reporters as biosensors. Growth of STEC (Sakai) transformed with any of the reporter plasmids (Supplementary Table 1) in defined medium with/out L-Ara*p* was not impaired compared to the vector only control (pKC026) (Supplementary Fig. 2). GFP levels from *araBA*_STEC_ (pACYC*araBA*_STEC_*::gfp+*, pJM058) increased in a dose-responsive manner to the concentration of L-Ara*p* when cultures were grown in either glycerol (R^2^=0.933, t=7.481, p=0.0017) or glucose (R^2^=0.973, t=12.012, p=0.0003) (Fig. 2). Expression of *araF*_STEC_ (pACYC*araF*_STEC_ ::*gfp*+, pJM066) increased in a similar manner (R^2^=0.891, t=4.674, p=0.0185), although expression was ∼1.5-fold lower than for *araBA*_STEC_ (Fig. 2). The specificity of the reporters was confirmed by growing STEC (Sakai) transformed with *araBA*_STEC_ (pJM058) or *araF*_STEC_ (pJM066) reporters in medium supplemented with xylose, which resulted in no expression of *araBA*_STEC_ (pACYC*araBAD*STEC*::gfp+*, pJM058). As expected *araF*_STEC_ (pACYC*araF*_STEC_::*gfp*+, pJM066) was induced in xylose as it is a low-affinity xylose transporter, but at a reduced level compared to arabinose (Supplementary Table 2). Utility of *araBA*_STEC_ (pACYC*araBAD*STEC*::gfp+*, pJM058) was also demonstrated during biofilm formation of non-pathogenic *E. coli* [32]. Therefore, the promoter constructs could effectively be used as specific biosensors for the presence of L-arabinose.

**Figure 2.**
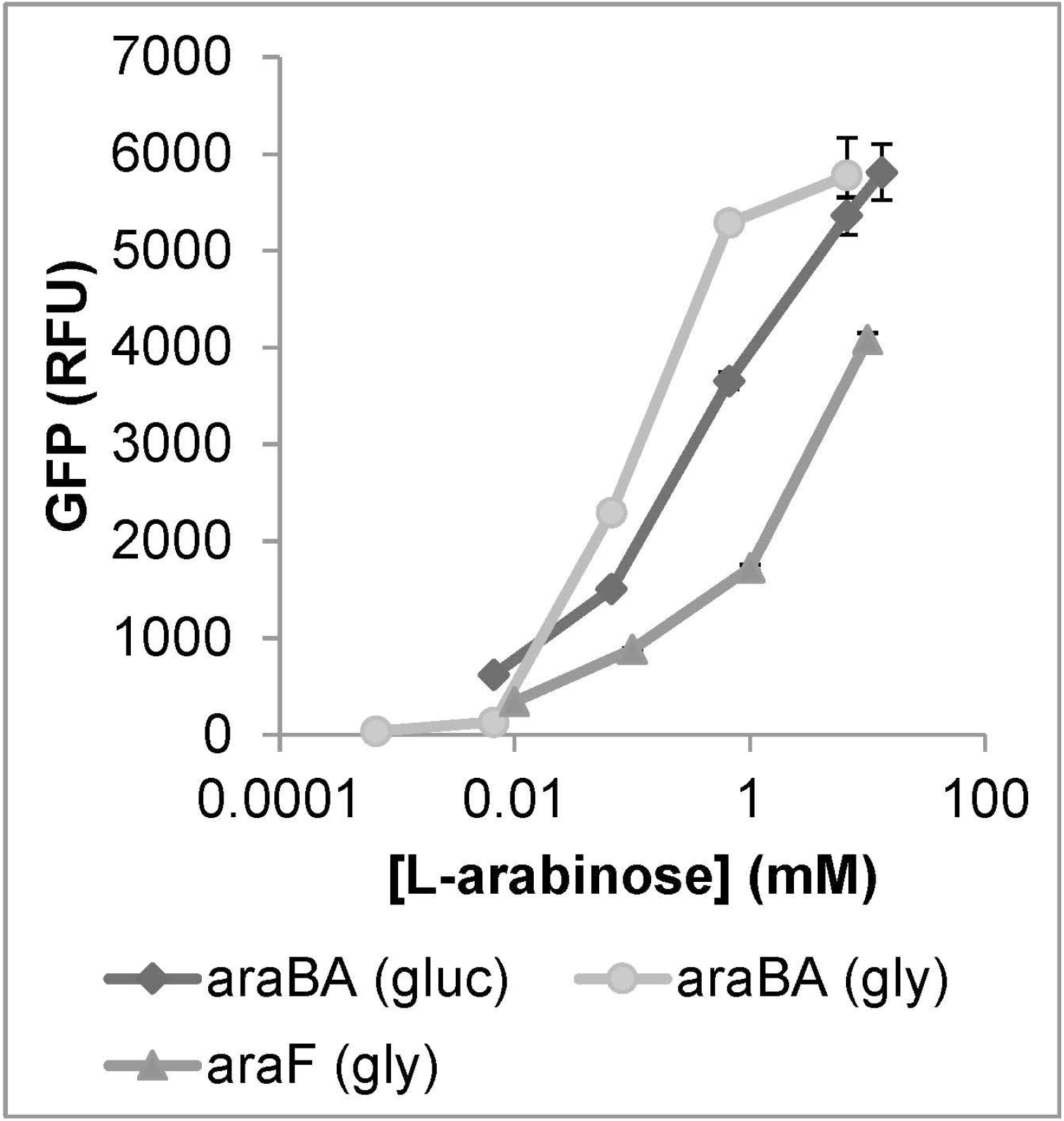
Dose-response expression of STEC (Sakai) *araBA::gfp+*. GFP measurements from STEC (Sakai) transformed with pACYC*araBA::gfp+* / pJM058 (*araBA*_STEC_) or pACYC*araF*_STEC_ ::*gfp+* / pJM066 (*araF*_Pba_) grown at 18 ° C in RD MOPS glucose (gluc) or RD MOPS glycerol (gly) supplemented with different concentrations of L-arabinose. Maximal GFP expression levels are presented from the mean of four replicates (+/- SD) at five h for pJM058 (*araBA*_STEC_) in glucose; eight h pJM058 (*araBA*_STEC_) in glycerol; or six h pJM066 (*araF*_Pba_) glycerol.

### Transcriptional activation of Pba (SCRI1043) *ara* genes *in vitro*

*P. atrosepticum* (Pba) and STEC O157:H7 isolates share a high proportion of their genomes [33] and since *ara* genes are widespread in bacteria, we hypothesised that L-arabinose metabolism and transport would be conserved between both species. The transcriptional activity of *araBA* and *araF* from Pba (strain 1043) were similarly quantified using the same GFP reporter system, in response to L-Ara*p*. Growth of Pba (1043) was prolonged at 18 °C in RD MOPS glycerol compared to STEC (Sakai), with a cell density of only 0.68 (OD_600_) after 24 h compared to ∼2.8 for STEC (Sakai), necessitating expression analysis to be carried out at 27 °C where they reached a similar densities (∼2.5). Although GFP expression was detectable from Pba (1043) transformed with either promoter fusion (pACYC*araBA*_Pba_::*gfp+*, pJM064; pACYC*araF*_Pba_::*gfp+* pJM065), growth was inhibited at > 1mM L-Ara*p* (Supplementary Fig. 2G). GFP was induced, but the levels exceeded linear limits of detection for the plate reader (> 600,000 RFU). Therefore, the promoters for *araBA* and *araF* were sub-cloned into the low copy number pSC101-based plasmid, with ∼ 6 copies / cell [34]. This strategy restored growth of the transformed Pba (1043) to that of the vector-only control (pWSK29) and the un-transformed strain (Supplementary Fig. 2G), even in the presence of 10 mM L-Ara*p*. The dose response of *araBA*_Pba_ (pJM067) and *araF*_Pba_ (pJM068) was similar to that of STEC (Sakai), and both Pba (1043) constructs produced maximal levels of GFP after 4 h in RD MOPS glycerol supplemented with 10 mM L-Ara*p*. Expression from the *araF*_Pba_ reporter was ∼ 2.5-fold higher than from the *araBA*_Pba_ reporter (Table 3). The response to xylose was similar to the STEC (Sakai) constructs, although the *araF*_Pba_ transporter reporter expression level in xylose was 40 % of the level in L-arabinose, showing some lack of specificity.

### Regulatory control of arabinose metabolism and transport in Pba and STEC

Genetic organisation of the *ara* loci are different between the species. In STEC arabinose metabolism and regulation linked and are *in cis*, while transport is *in trans* and dependent on AraC (and CRP) binding. In Pba (1043) metabolism, transport and regulation are linked *in cis*, but the epimerase is located *in trans*, and *araC* is embedded within the transport polycistronic operon at the 3’-end (Supplemental Fig. 3A). The STEC *araBA* genes are co-transcribed and the epimerase, *araD* is adjacent but assumed to be transcribed separately via a canonical ribosome binding site, while *araC* is divergently transcribed from *araBA* [35]. These distinct genetic organisations appear to be conserved is the wider genera of both species. Six REP sequences are present 5’ to the *araD*_STEC_ CDS. The AraC regulatory protein sequences share 57 % overall amino acid identity, with conservation in all L-Ara*p* binding residues, 8/10 beta barrels, one of the dimerisation alpha helices and the DNA- binding / H-T-H domain (Supplemental Fig. 3B)

To compare the STEC and Pba L-arabinose regulatory networks, exchange experiments were performed, where the L-arabinose biosensor plasmids were transformed into the non-native host. This artificial situation was used for a direct comparison between the species. Growth of the pACYC-based Pba reporters transformed into STEC (Sakai) was not inhibited in the presence of L-Ara*p*, allowing direct comparison. For each pair of strain backgrounds, expression was highest in Pba (1043), at levels up to 8.6-fold for the *araF*_Pba_ reporter (Table 3). The least amount of difference was for *araBAD*_STEC_ where the levels of expression were relatively similar, but still higher in the Pba (1043) background. There was a difference between each pair of *ara* operons, such that the *araF*_Pba_ promoter was stronger than *araBA*_Pba_, whereas the opposite was true for the STEC (Sakai). Furthermore, *araF*_STEC_ was over-induced in the Pba (1043) background, reversing the trend seen in the native STEC background. Hence, in Pba (1043), arabinose transport via the high affinity AraF transporter is particularly highly induced, even when the *araF* promoter is non-native, which is the opposite of the situation observed in STEC.

Regulatory control via AraC was examined with an *araC* deletion mutant for STEC (Sakai). Under *in vitro* conditions, the *araBAD*_STEC_ reporter construct was still induced in this background, to levels marginally higher than the WT (Table 3), (as expected due to negative feedback regulation of *araBA*) whereas *araF*_STEC_ was completely repressed in the absence of *araC*. In contrast, the Pba reporter *araBA*_Pba_ was repressed in the absence of *araC* in the STEC (Sakai) background, whereas *araF*_Pba_ was still expressed to ∼ 50 % of that in the WT STEC (Sakai) background (Table 3). Taken together, the data shows differences in regulatory control and potentially specificity in AraC-dependent regulation, with some level of constitutive expression of *araF*_Pba_.

### Expression of ara genes in plant extracts and in planta

Since L-arabinose is normally complexed in plant tissue as L-Ara*f, in vitro* plant extracts were used for quantification of gene expression in the absence of other factors. Expression from STEC (Sakai) transformed with the *araBA*_STEC_ reporter was assessed *in vitro* in media supplemented with a range of different plant extracts, to determine whether L-arabinose dependent-activity from the promoters could be quantified. Detection of *araBA*_STEC_ was absent (spinach, lettuce) or minimal (tomato) over five days of exposure in the plant extracts compared to the medium with L-Ara*p* added (Table 2), which indicated that the levels of L-arabinose were either below 0.6 µM or components in the plant extracts masked GFP detection by the plate reader. Therefore, a qualitative approach using microscopy was taken to determine if *ara* gene expression could be detected *in planta* over longer time periods.

**Table 2.** GFP measurements from *araBAD*_STEC_ in STEC (Sakai) inoculated with different plant extracts types.

| Medium | Day 0 | Day 1 | Day 2 | Day 5 |
| --- | --- | --- | --- | --- |
| Spinach Leaf Lysate | <0 | <0 | <0 | \$ |
| Tomato Leaf Lysate | 243.600 | <0 | 1,494.266 | <0 |
| Lettuce Leaf Lysate | <0 | <0 | <0 | <0 |
| RDM L-ara 6.661 mM | 14.293 | 20,234.293 | 58,610.325 | 64,072.291 |
| RDM L-ara 0.666 mM | 10.193 | 10,329.918 | 24,307.419 | 28,889.353 |
| RDM L-ara 66.609 µM | 19.933 | 5,752.784 | 1,342.746 | 1,733.018 |
| RDM L-ara 6.661 µM | 29.340 | 576.825 | 292.451 | 415.993 |
| RDM L-ara 0.666 µM | 17.607 | <0 | 45.755 | 330.483 |
| RD MOPS only | 30.287 | 42.179 | <0 | <0 |
| RDM D-glucose 0.2% | 38.473 | 23.344 | <0 | 99.814 |

\* values that were less than 0 after normalisation are indicated by '<0'; \$, the cells coagulated

Expression was assessed qualitatively from bacteria inoculated into the internal tissue of leaves and from leaf surfaces. Two plant models were chosen that are known to support high levels of colonisation and in an attempt to override differences in the plant-microbe interactions, i.e. potential PCWDE production by Pba. *N. benthamiana* was used for infiltrated bacteria, since unlike spinach or lettuce, it allows unrestricted growth of internalised *E. coli* O157:H7 (Sakai) and Pba (1043) [36]; surface-associated bacteria were assessed from inoculation of broccoli microgreens, as a representative edible crop species that supports high growth of both isolates [29]. Colonies formed in both species after 11 days in the *N. benthamiana* apoplast, and they both expressed metabolism, *araBA* and transport, *araF* genes in native backgrounds (Fig. 3a-d). Some variation in expression was evident at the single cell level in all four constructs. Both STEC and Pba formed extensive colonies on the plant cell wall margins of broccoli cotyledons after 6 days, and both expressed metabolism and transport reporter genes (Fig. 3 e-h). Variation at the single cell level was evident (quantification on plant leaves was not possible). No GFP was detected from the empty vector (pKC026) from either STEC or Pba in broccoli or *N. benthamiana* (Suppl. Fig 4a-b). Thus, both species expressed L- ararbinose metabolism and transport genes *in planta*, under different model systems.

**Figure 3.**
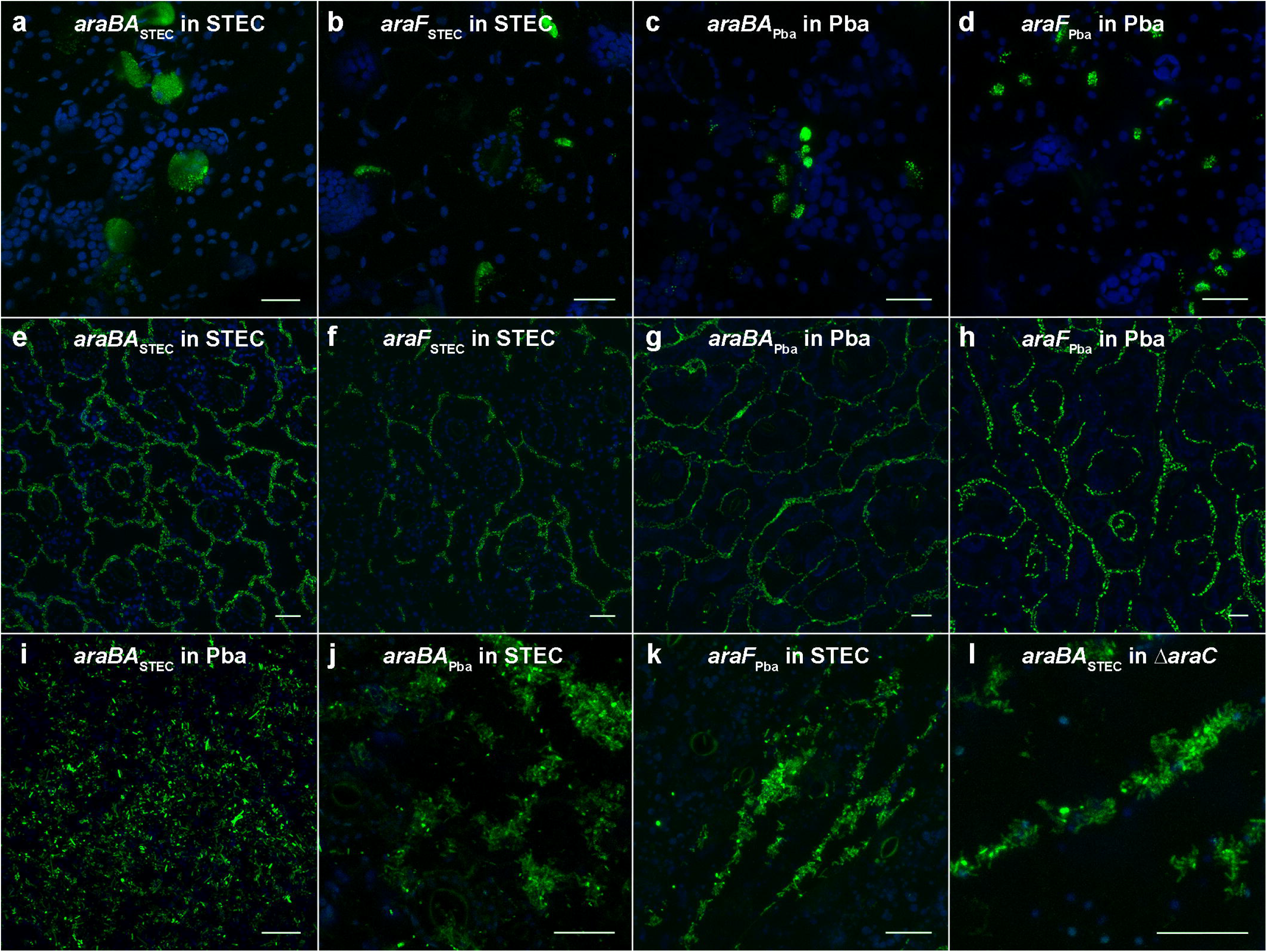
*In planta* expression of arabinose metabolism and transport genes. GFP reporter expression of *araBA* and *araF* plasmid constructs in STEC (Sakai), Pba (1043) or STEC (Sakai Δ*araC*) backgrounds. Reporter constructs transformed in native backgrounds were examined from bacteria infiltrated into *N. benthamiana* leaves and colonies imaged at day 11 (a), day 4 (b, c) or day 1 (d) after inoculation. Reporter constructs transformed in native or non-native backgrounds were examined on broccoli cotyledons germinated from inoculated (imbibed) seeds. Bacterial colonies of STEC (e, f, j, and k), Pba (g, h, and i) or Δ*araC* (l) were present on the surfaces of cotyledons at day 5 (l), day 6 (e, j, k), day 7 (i), day 10 (g, h) and day 11 (f). Chloroplasts in the epidermal and mesophyll cells are shown (blue). All scale bars represent 25 µm.

To see whether *ara* gene regulatory control observed *in vitro* held *in planta*, expression of *araBA*_STEC_ (pJM058) transformed into Pba (1043) and *araBA*_Pba_ (pJM064) transformed into STEC was examined from surface-associated bacteria on broccoli leaves. Both constructs were expressed *in planta* in the non-native backgrounds, although morphological differences occurred in Pba cells presumably due to GFP toxicity (Fig. 3 i-k). Variation in expression at the single cell level was evident and in STEC: GFP levels from most cells was low with only a small number expressing a high level. A similar pattern occurred for STEC hosting *araF*_Pba_. GFP was only detected from the *araBA*_STEC_ reporter in the STEC Δ*araC* mutant background (Fig. 3l), in keeping with quantitative data (Table 3). There was no detectable GFP from *araF*_STEC_ or either of the Pba contructs *araBA*_Pba_ or *araF*_Pba_ in this background (Suppl. Fig. 4d-f), confirming their responsiveness to L-arabinose.

Since Pba (1043) encodes PCWDE, the responsiveness of the *araBA*_Pba_ reporter was quantified *in vitro* from incubation with a crude preparation of leaf cell wall polysaccharides (spinach). Glycerol was added as a carbon source to facilitate bacterial growth and increased glycerol concentration used to promote induction of any PCWDE [37], although their expression was not measured *per se*. Growth of Pba (1043) reached comparable densities in all substrates and was not affected by variable glycerol concentration. GFP levels for *araBA*_Pba_ reached ∼ 450 RFU in the presence of cell wall polysaccharide extract with 0.4 % glycerol, compared to ∼150 RFU in their absence, with no difference between 6- and 24-h incubation (Fig. 4a). This level of expression was ∼ 10-fold lower compared to *in vitro* response to L-Ara*p* (Table 3), indicative of the difference in response to free L- Ara*p* and complexed L-Ara*f*. In contrast, no expression was detected from the STEC (Sakai) *araBA*_STEC_ reporter in cell wall polysaccharides. However, incubation with purified L-Ara*f* induced expression of STEC (Sakai) metabolism genes *araA* and *araD* in the presence the debranched form of pectin or as oligomers as arabinobiose and arabinoheptose, while the fully branched form of pectin did not (Fig. 4b). Furthermore, STEEC (Sakai) showed marginal growth in minimal media supplemented with just arabinobiose or arabinoheptose, reaching 6.716 (± 0.240) and 7.024 (± 0.724) Log_10_ cfu/ml (respectively) after 48 h at 18 °C from an inoculum of 6 Log_10_ cfu/ml, compared with 8.643 (± 0.058) or 8.134 (± 0.365) Log_10_ cfu/ml in medium supplemented with L-Ara*p* or glucose (respectively). No growth occurred in a BSA-supplemented control medium. Therefore, STEC (Sakai) was responsive to L-Ara*f* in oligosaccharides *in vitro* and able to utilise them for growth.

**Table 3.** Expression of *ara* genes in native and different backgrounds.

| Plasmid / Isolate | STEC (Sakai) | Pba (1043) | STEC Δ*araC* (Sakai) |
| --- | --- | --- | --- |
| pACYC*araBAD*<sub>STEC</sub>::*gfp*+ (pJM058) | 11,151 (± 493) | 15,566 (± 93) | 15,809 (± 224) |
| pACYC*araF*<sub>STEC</sub>::*gfp*+ (pJM066) | 4,086 (± 64) | 35,215 (± 560) | 275 (± 3) |
| pWSK*araBA*<sub>Pba</sub>::*gfp*+ (pJM067) | 1,882 (± 138) | 10,213 (± 182) | NT |
| pWSK*araF*<sub>Pba</sub>::*gfp*+ (pJM068) | 3,278 (± 260) | 25,075 (± 1804) | NT |
| pACYC*araBA*<sub>Pba</sub>::*gfp*+ (pJM064) | 5,050 (± 124) | * | 186 (± 3) |
| pACYC*araF*<sub>Pba</sub>::*gfp*+ (pJM065) | 4,868 (± 81) ** | * | 1,747 (± 20) ** |

(representative experimental replicate, total 4 reps)
NT: not tested
\* > 600,000 RFU
\*\* high background: 1,100 – 1,300 RFU in the absence of arabinose
GFP was measured in response to 10 mM L-arabinose at maximal expression times (four or six h), for
cultures grown in RD MOPS glycerol at 27 °C. Blue background indicates native expression and red
indicates non-host expression.

**Figure 4.**
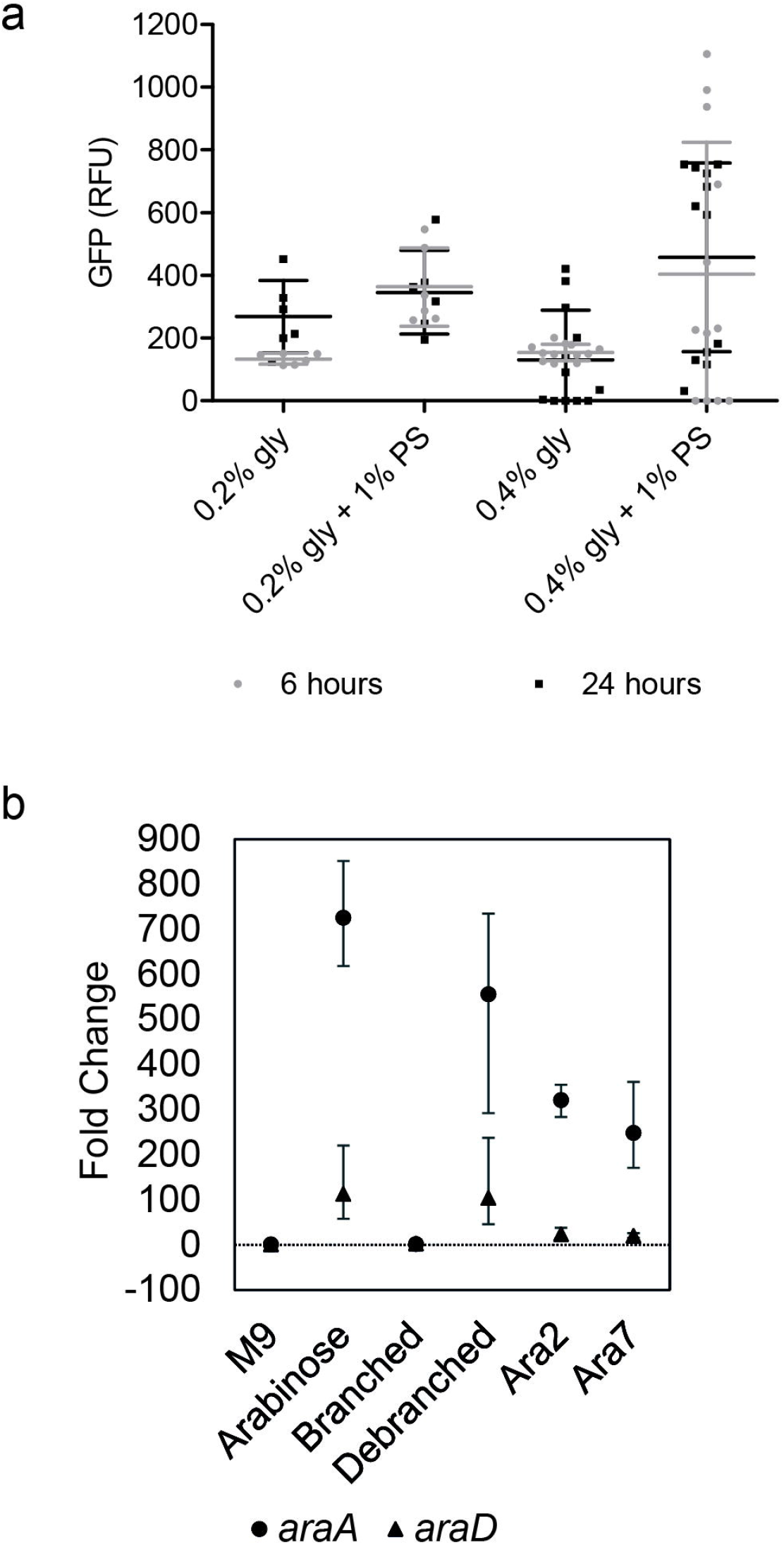
Expression of *araBA*_Pba_ in the presence of plant cell wall polysaccharides. Reporter expression for *araBA*_Pba_ (pJM067, pWSK*araBA*_Pba_*::gfp+*) in Pba (1043) following incubation in 1% (w/v) spinach alcohol-insoluble polysaccharide (1 % PS) extract at two glycerol concentrations (0.2, 0.4 %), for 6 or 24 h at 27 °C (a). Fluorescence measurements, normalised for cell density, are expressed as average RFU (+/- SD) relative to the control plasmid, from triplicate samples x two or four independent experiments (for 0.2 % or 0.4 % glycerol, respectively). Gene expression measured directly from STEC (Sakai) *araA* and *araD* (b) after one h in M9 minimal medium only (M9); or supplemented with 0.2 % L-arabinose (Arabinose); (1-5)-α-linked pectin backbone which contains (1- 3)-α-linked and possibly (1,2)-α-linked L-arabinofuranosyl residues (Branched); (1-5)-α-linked backbone treated with α-L-arabinofuranosidase (Debranched); (1-5)-α-L-arabinobiose (Ara2); and (1,5)-α-L-arabinoheptose (Ara7). Gene expression is expressed as averaged fold-change (+/- SD) relative to the reference gene (*gyrB*), from three biological x three technical replicates.

### Arabinan complement *in planta*

L-arabinose is abundant present in plant cell wall pectin and glycoproteins as L-Ara*f*, while L-Ara*p* is much less so and a minor glycan in plants in comparison to glucose, sucrose and fructose [4]. In crude leaf cell wall polysaccharide extracts of spinach and lettuce, spinach contained five-fold more L-Ara*p* compared to lettuce leaf extracts [16]. A screen of pectin-enriched fractions (CDTA treatment) from 15 plant horticultural species and cultivars, with different tissues (leaves, root or sprouted seeds) with antibody probes confirmed that arabinans as L-Ara*f* were more abundant of spinach than in lettuce across a range of cultivars [27]. It also showed that broccoli microgreens were comparable to sprouted seeds for linear pentasaccharides of L-Ara*f* ((1-5)-α-L-arabinan, LM6) (Supplemental Table 3). Closer examination of broccoli microgreen-leaf showed (1-5)-α-L-arabinan (LM6) was dominant in the pectin-enriched fraction together with homogalacturonan (JIM7), and AGP (LM2) as the dominant glycoprotein (Fig. 5a). Processed arabinans in RG-I were detected in broccoli as a minor component (LM16). NaOH-derived enrichments detected xyloglucans as a principle component of hemicellulose (LM15, LM25). The distribution of (1-5)-α-L-arabinan (LM6) was visualised in broccoli cotyledons by antibody staining (Fig. 5b), with or without inoculated STEC (Sakai). (1-5)-α-L-arabinan was distributed along cell margins (LM6, green) and coincided with STEC (Sakai) localisation (α-O157 antibody, magenta).

**Figure 5.**
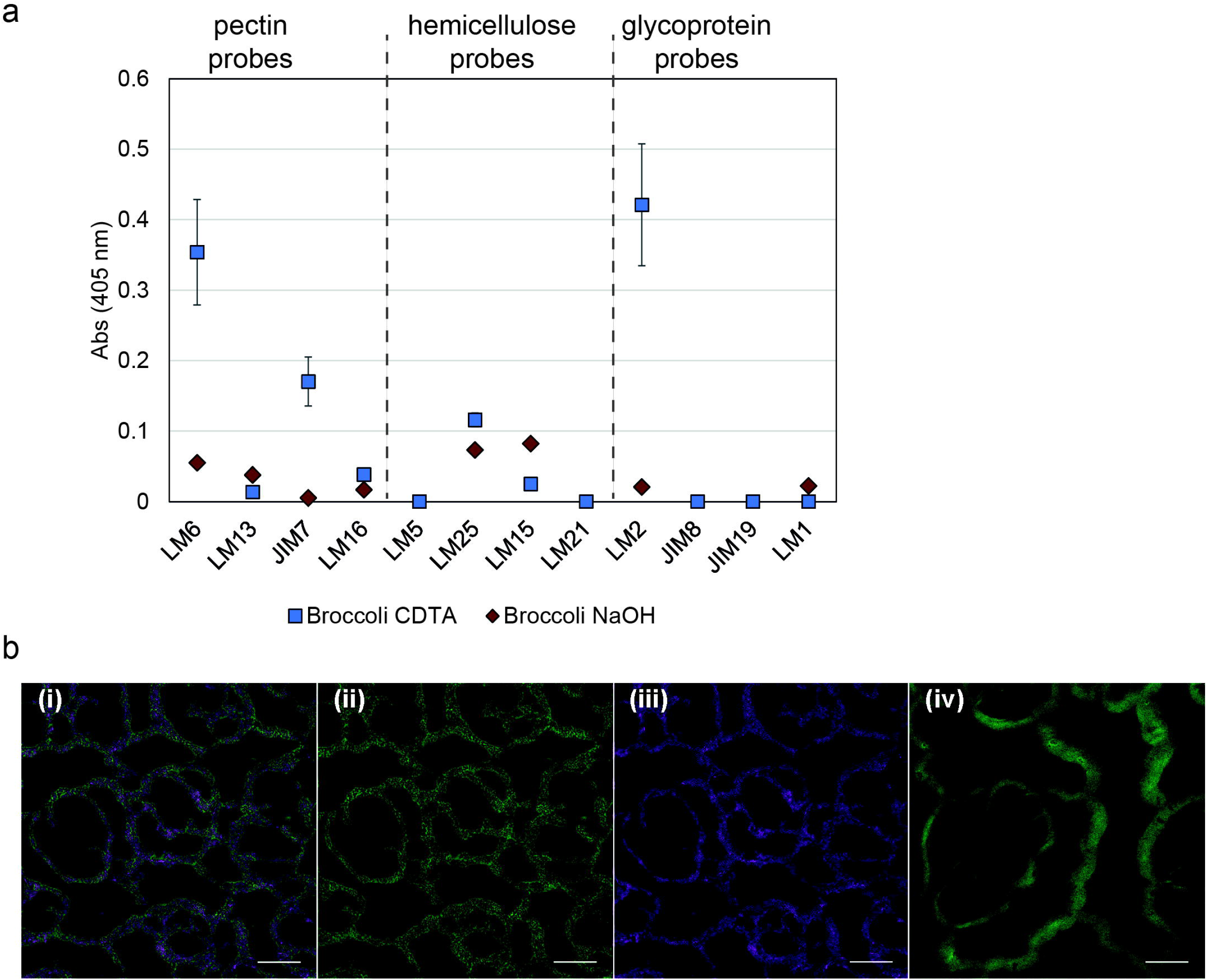
Glycan complement of broccoli in plant cell wall polysaccharide extracts. Alcohol insoluble residues for broccoli microgreen enriched for pectin by CDTA extraction (blue) or hemicellulose by NaOH extraction (brown) (a). ELISA quantification for indicated primary antibodies probes with secondary antibody conjugated with HRP, is shown as absorbance at 405 nm, averaged (+/- SD) from four extractions x two technical repeats. One-way ANOVA showed significant differences between antibodies for both extractions (pectin P = 1.11 x 10 ^-4^; hemicelluose P = 0.043). Detection of (1-5)-α-L-arabinans in broccoli cotyledon tissue (b). Broccoli cotyledons, seven days after germination in the presence of *E. coli* STEC Sakai (WT), treated with LM6 primary antibody, labelled with Alexa Fluor^®^ 488 nm (green) and α-O157 primary antibody, labelled with Alexa Fluor^®^ 568 nm (magenta) secondary antibody showing single-layer confocal images of inoculated (i, ii, iii) or non-inoculated tissue (iv). All scale bars represent 25 µm.

## Discussion

In STEC (Sakai), L-arabinose metabolism is active in plants but non-essential for its colonisation: *ara* genes were responsive *in vitro* to leaf extracts and *in planta* during colonisation for a range of plant models, but the gene cluster knock-out was not affected in spinach or lettuce colonisation over a 10- day period. Therefore, it appears that STEC (Sakai) is able to take advantage of L-arabinose as an abundant plant-derived metabolite in an opportunistic manner. Use of different plant models to demonstrate either surface or internal plant colonisation showed that STEC (Sakai) responded to L-arabinose once it establishes *in planta* colonies. Repression of metabolic genes on initial introduction into plants suggests that a period of adaption occurs before the L-arabinose system is induced *in planta*. The lack of dependency on L-arabinose metabolism by STEC (Sakai) in plant colonisation is indicative of metabolic redundancy, in line with the non-essentiality of the *ara* gene cluster for growth in rich media [38], and in keeping with *E. coli* as a heterotroph [39].

For microbes that cross biological kingdoms, metabolic diversity is an essential asset. The *Enterobacteriaceae* tend to have mosaic genomes contributing to such generalism, which has also been described for plant or animal-associated pseudomonads [40]. Intestinal microbiota within animal hosts can access and metabolise ingested plant material. For example, activity of *E. coli* KduI facilitates conversion of pectin-derived D-galacturonate in gnotobiotic mice [41]. In nature, fermentation of hexoses occurs through the Entner Duodoroff (ED) pathway, which we found was induced in STEC (Sakai) on *in vitro* exposure to plant extracts [16]. Discovery of non-phosphorylative pathway for pentoses, including L-arabinose, may also facilitate colonisation of plant niches as it has an equivalent role to the ED pathway [42]. Furthermore, promiscuity in pentose transporters broadens metabolite options when gylcans become scare [43], e.g. on leaf surfaces.

Examination of a range of plant species showed that arabinans (as L-Ara*f*) were differently distributed in the pectin and AGP components between plant species and tissues. L-Ara*p* is present in leaf cell wall polysaccharides at relatively low levels compared to other glycans but also with still plant-species and tissue differences [4, 16]. STEC (Sakai) is responsive to L-Ara*f* in debranched pectin and as arabinan oligomers as well as L-Ara*p*, but not in branched pectin. We previously showed that STEC (Sakai) ECP targeted (1-5)-α-L-arabinans present in RG1 of pectin for adherence to plant cells [27], which may account in part at least, for observed differences in colonisation. This raises a potential link between fimbrial-dependent attachment and metabolism to facilitate plant colonisation, as has been seen in uropathogenic *E. coli* with type 1 fimbriae [39]. Since *E. coli* do not encode pectinases and cannot access L-Ara*f* directly, there are two potential sources for STEC *in planta*: from turn-over of cell wall components, e.g. during plant growth and development from activity of plant-derived α-L-arabinofuranosidase glycoside hydrolases [3], and / or from pectinase activities of the endemic microbiome [44]. All plant inoculation experiments for STEC (Sakai) included endemic microbiomes in our work incurring microbial ecological interactions, whether competitive, neutral or mutualistic. As such, for STEC to have sufficient ecological fitness to invade established microbiomes, capacity for diverse metabolite utilisation is critical.

*P. atrosepticum* is a member of the *Enterobacteriaceae* that differs from *E. coli* in that it encodes PCWDE, making them a useful comparison pair for metabolite use. Pba L-arabinose metabolism appears to be linked to PCWDE expression through the response regulator ExpM (RssB in *E. coli*) [45] since *araA, C* and *H* were significantly induced > 2-fold after 12 h inoculation on potato tubers in a Pba *expM* mutant [22]. The quorum-sensing dependent RsmA-*rsmB* negative regulatory system (CsrA-*csrB* in *E. coli*) links metabolism to production and secretion of PCWDE through RpoS as a transcriptional activator of the RsmA [22]. ExpM/RssB negatively regulates RpoS that when phosphorylated, facilitates degradation of the sigma factor by ClpXP protease activity, and phosphorylation of ExpM / RssB is controlled (in part) by the sensor kinase ArcB, which is itself inactivated during carbon starvation [46]. Hence ExpM activity alleviates RsmA-dependent repression of the PCWDE. Therefore, the elevated RpoS levels that would occur in the *expM* mutant or in the presence of inactive, unphosphorylated ExpM induce expression of alternative metabolic pathways, including L-arabinose. The ExpM-dependent effect appeared specific for arabinose since the xylose genes were not similarly induced in Pba, and the *ara* genes were not impacted by other key virulence regulator mutants [22, 47].

Despite the taxonomic similarity of STEC (Sakai) and Pba (1043), differences exist in the genetic organisation, expression profiles and regulation of L-arabinose metabolism and transport. L-arabinose transport appears to be more sensitive in Pba compared to STEC, since *in vitro* expression of *araF* was ∼ 3-fold higher than the metabolic genes *araBA*, whereas the converse occurred in STEC. There was also evidence for constitutive, or non-AraC dependent expression of Pba L-arabinose transport (via AraF), whereas STEC *araF* induction was completely dependent on AraC. Although the AraC amino acid sequences for L-arabinose binding and DNA-binding domains were conserved in both species, there is sufficient species-dependent specificity such that negative feedback of *araBA* by AraC was lost when the *Pba* genes were expressed in the STEC *araC* mutant, and *araF*_STEC_ was over-induced in the Pba background. Although it was not possible to make direct comparisons between Pba and STEC in *in planta*, expression of the biosensor reporters echoed the *in vitro* expression patterns for both species, thus provided a faithful representation.

STEC and Pba have different primary reservoirs (ruminants and potatoes, respectively), and as such, STEC can be considered an invader of plant niches [48]. Within the same environment e.g. plants niches, different traits will contribute to their relative fitness. Although both species possess metabolic flexibility, nuanced regulatory differences and the link to PCWDE expression account for their different strategies for L-arabinose metabolism. In STEC, the system is primed toward opportunistic access, weighted to catabolic activation, while in Pba, the system is primed towards direct access through high-affinity transport. This has enabled STEC to exploit such an abundant plant metabolite during colonisation of plants, but not in a dependent manner.

## Acknowledgements

NH, RWJ, IT & LC were supported by a PhD studentship funded by the James Hutton Institute and the University of Reading; NH, JM, AH, BM, SH & KW were supported by the Scottish Government Strategic funding (RD3.1.3, RD2.3.3); NH & YR was supported by a Leverhulme Trust grant (RPG-096). We are grateful for support of Prof. Simon Andrews and Prof. Carol Wagstaff (University of Reading) to LC during her PhD. The *araC* knockout mutant was a generous gift from Prof. Andrew Roe, University of Glasgow, constructed by Dr James Connolly (currently University of Newcastle).

## Supplemental Table & Figure Legends

**Supplemental Figure 1 Expression of E. coli O157:H7 araA in spinach and lettuce extracts**

Expression of *araA* quantified by the DNA microarray, or by qPCR from the microarray samples (array) and independent repeated samples (repeat) in plant extracts (a), or following infiltration into leaves for one h (b). The culture controls were bacteria incubated in no-plant, vermiculite control, or in in the MgCl_2_ infiltration medium. The average of three biological replicates, each with three technical replicates, with the SD.

**Supplementary Figure 2 Growth curves of STEC (Sakai) and Pba (1043) transformed with fluorescent reporters**

Fluorescent reporters pJM058 (A, B), pJM064 (C, D, G), pKC026 (E, F, G), pJM065, pJM067, pJM068 or pWSK029 (G) were transformed into STEC (Sakai WT) (A, C, E), STEC (Sakai Δ*araC*) (B, D, F), or Pba (1043) (G). Expression was monitored over time at 27 °C at arabinose concentrations of 0, 0.01, 0.1, 1 or 10 mM as indicated (A-F), or just at 10 mM (G).

**Supplementary Figure 3 Genetic and structural organisation of the ara loci for STEC (Sakai) and Pba (1043)**.

Organisation of the genetic loci (a), with the metabolism genes in blue (*araB*, ribulokinase; *araA*, isomerase; *araD*, epimerase); transport genes in green (*araF*, arabinose binding protein; *araG*, ATP symporer; *araH*, permease); and regulator in orange (*araC*, binds L-arabinose). Gene clusters or genes that are in *cis* are marked by ‘//’. The STEC (Sakai) *araFGH* is on the complementary strand on the chromosome, and the CDS for *araH* is annotated as split between ECs2607 and ECs2606. Genetic distances are approximated on the gene clusters. Alignment of the AraC CDS (b) indicate the location of secondary structure features (as annotated for STEC Sakai).

**Supplementary Figure 4 *In planta* detection of GFP for controls and non- expressing constructs**

The empty GFP reporter vector, pKC026, was transformed in STEC (Sakai) or Pba (1043) to assess any background fluorescence. Epidermal and mesophyll cells of a leaf of *Nicotiana benthamiana* 4 days after infiltration with empty vector control pKC026 in Pba (a). Inoculated (seed imbibed) broccoli was harvested 6 days after germination for detection of plasmid-containing bacteria, treated with α- O157 primary antibody and labelled with Alexa Fluor^®^ 568 nm secondary antibody (magenta) for detection on non-GFP STEC (Sakai). STEC WT transformed with pKC026 (b) or STEC Δ*araC* transformed *araF*_STEC_, *araBA*_Pba_ and *araF*_Pba_ constructs (c- f) were imaged on the epidermis with chloroplasts (blue) indicating the position of the epidermal and mesophyll cells. All scale bars represent 25 µm.

