## Supplementary figures and images for "The role of L-arabinose metabolism for *Escherichia coli* O157:H7 in edible plants"

### Supplementary Figure 1

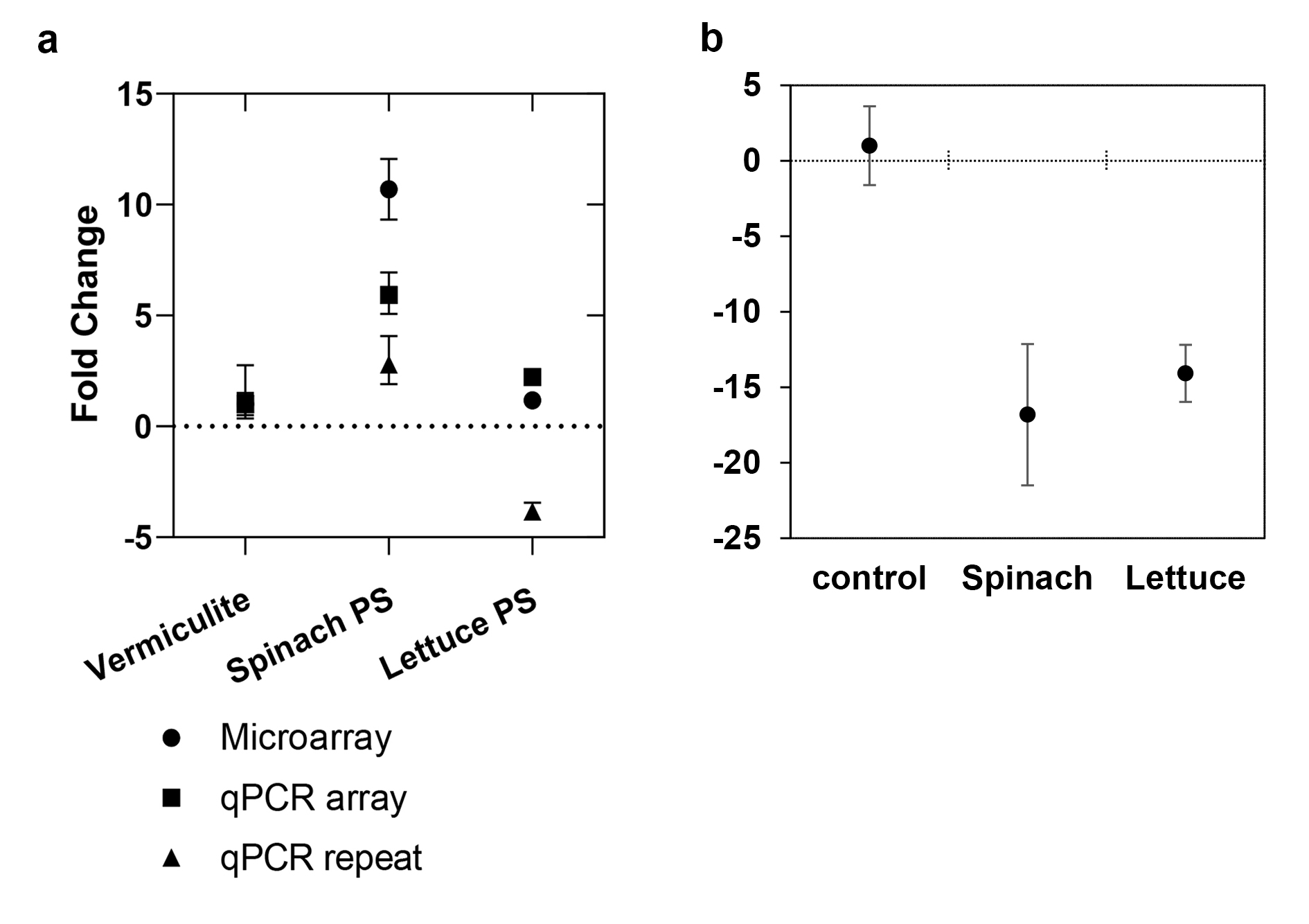

### Supplementary Figure 2

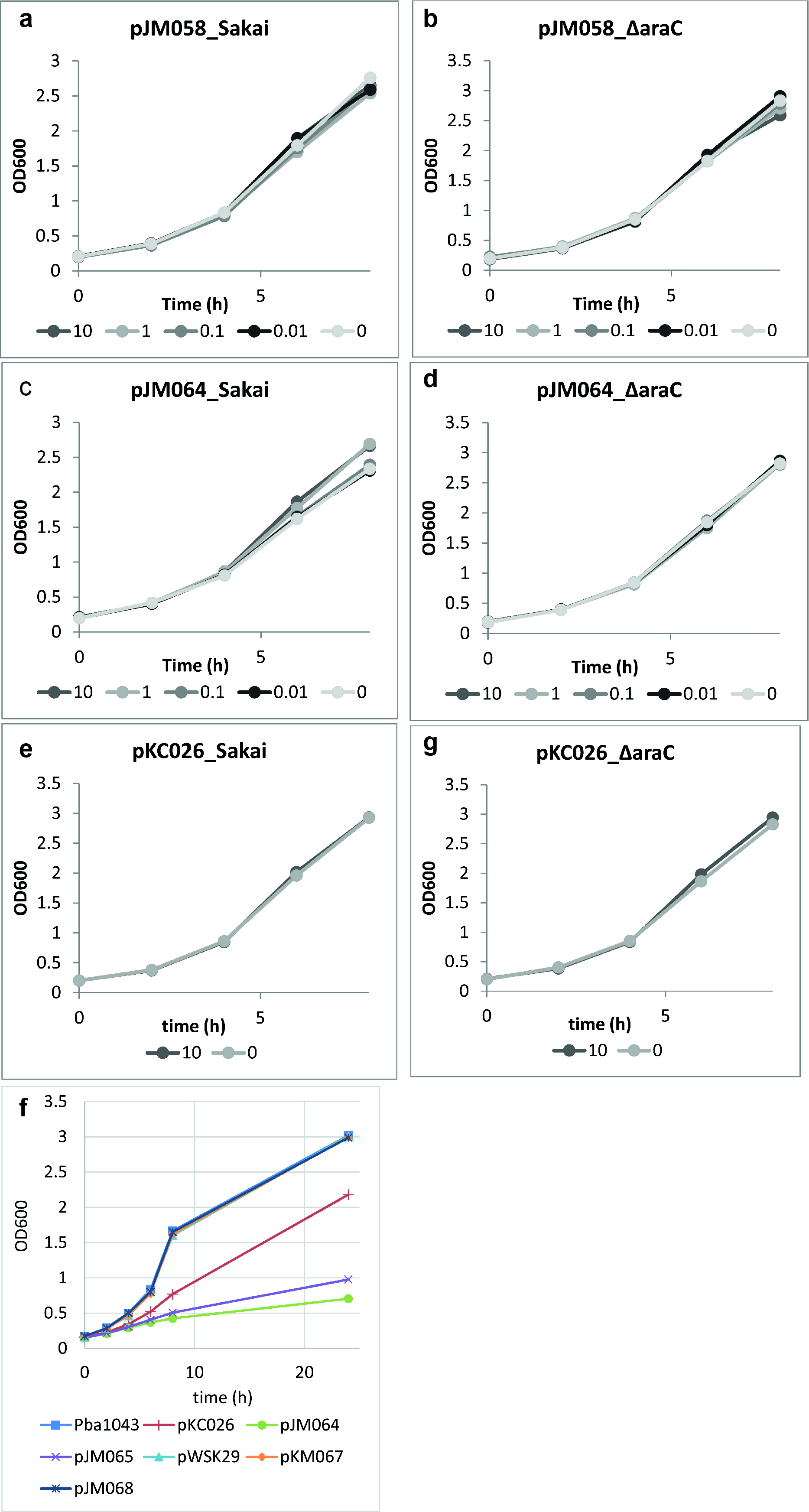

### Supplementary Figure 3

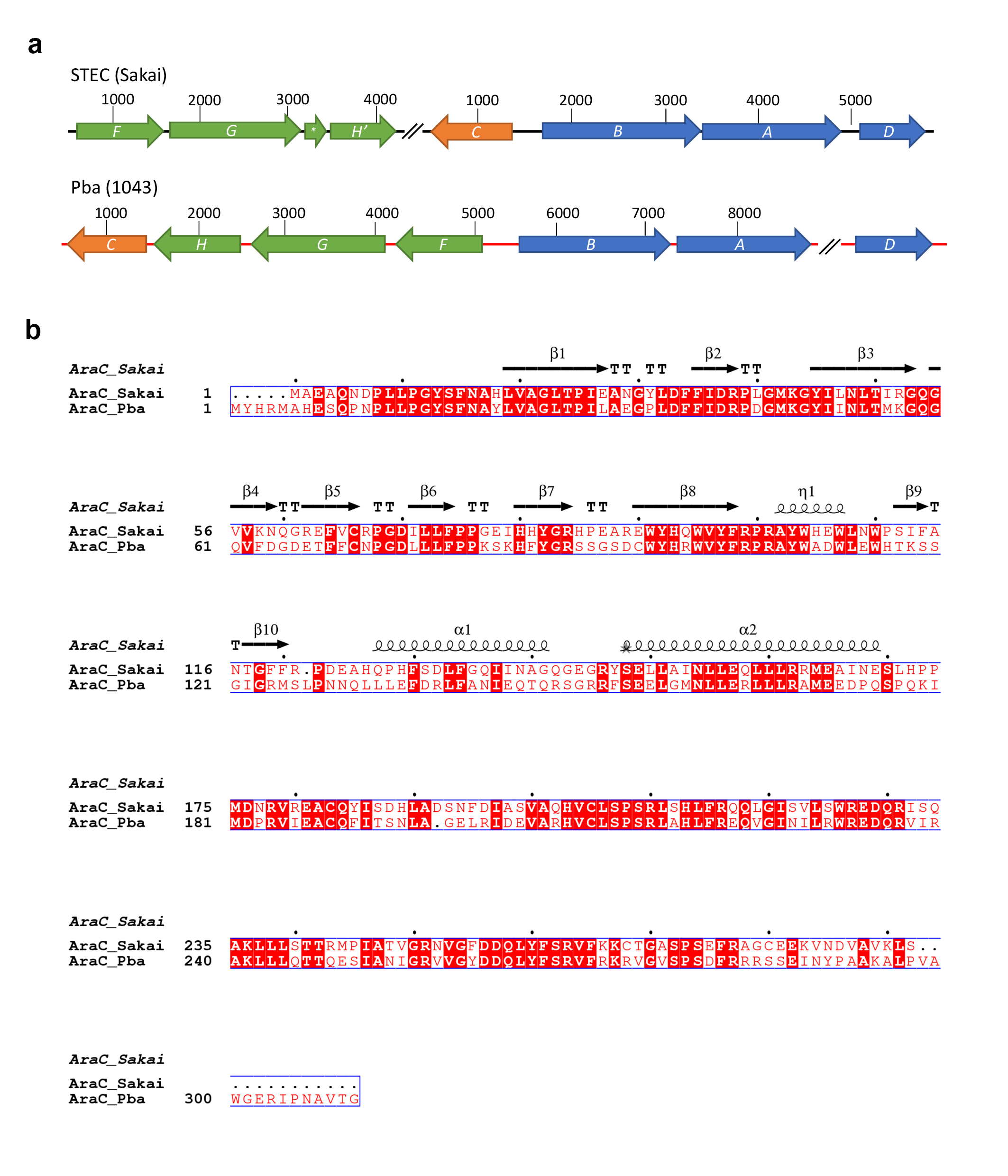

### Supplementary Figure 4

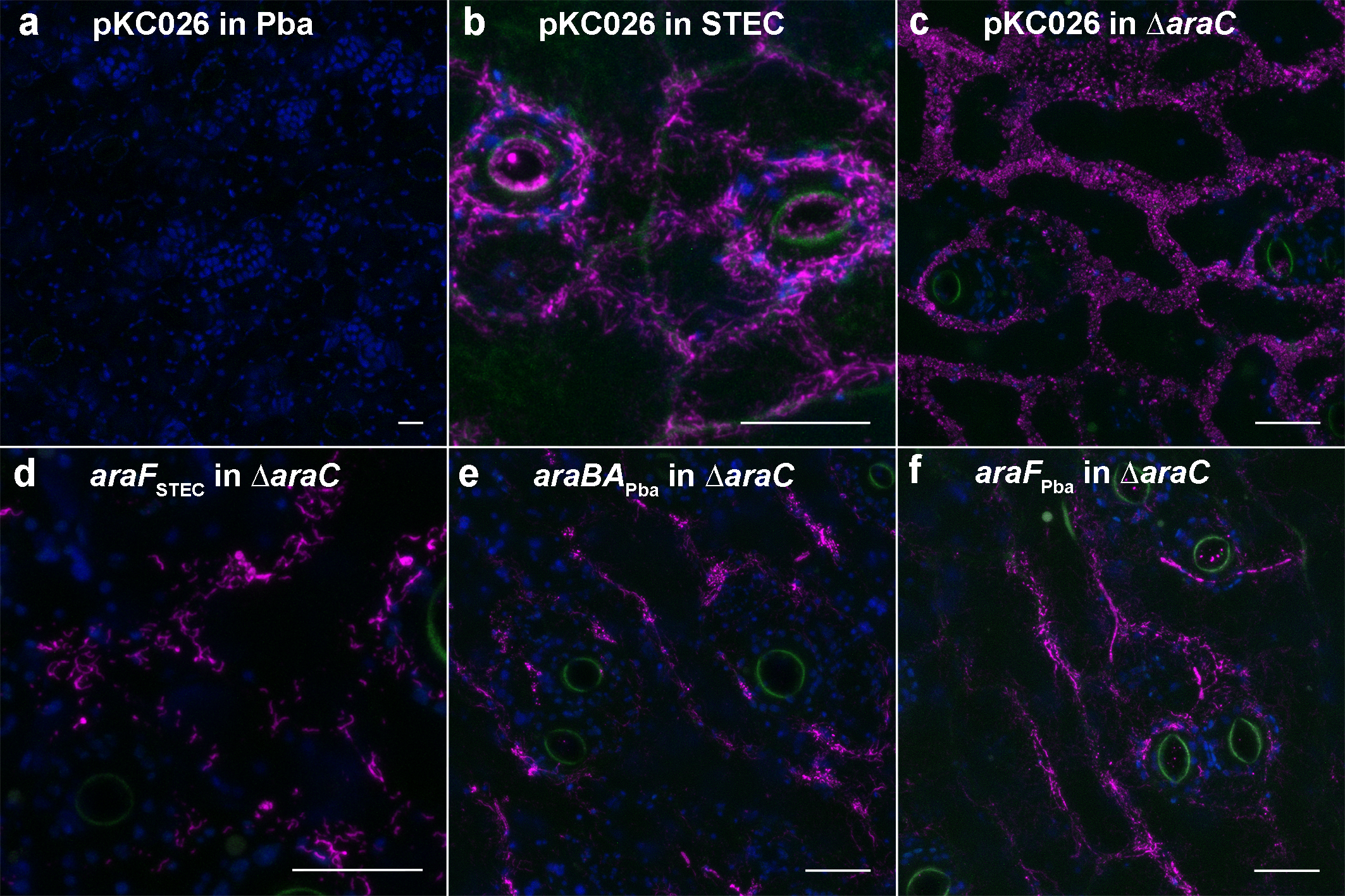
