## Supplementary Tables for "The role of L-arabinose metabolism for *Escherichia coli* O157:H7 in edible plants"

**Supplemental Data**

**Supplementary Table 1	Plasmids and primers used in the study**

| **Plasmid /**  **genotype** | **Gene / accession #** | **Vector** | **Cloning primers** |
| --- | --- | --- | --- |
| pJM058  pACYC*araBAD_STEC_::gfp+* | STEC (Sakai) *araBAD*  ECs0067, ECs0066, ECs0065 | pKC026 | 5’-GGTCTAGAGCCTGTCAAATGGACGAAGC  5’-CAGGTCGACTCTAGAGGATCC |
| pJM066  pACYC*araF_STEC_::gfp+* | STEC (Sakai) *araF*  ECs2609 | pKC026 | 5’-GCTCTAGAGATGGCTCTCATTATACG  5’-CTCTAGAGCTTTAGTGTCGTTTTGTGCAGG |
| pJM064  pACYC*araBA_Pba_::gfp+* | Pba (SRCI1043) *araBA* ECARS11170, ECARS11175 | pKC026 | 5’-GCTCTAGAGGTGTAATTATCTTTTCATTCGC  5’-GCTCTAGACTCATTAATCTCAGATCGTTGG |
| pJM065  pACYC*araF_Pba_::gfp+* | Pba (SRCI1043) *araF*  ECORS11165 | pKC026 | 5’-GCTCTAGACTCATTAATCTCAGATCGTTGG  5’-GCTCTAGAGGTGTAATTATCTTTTCATTCGC |
| pJM067  pWSK*araBA_Pba_::gfp+* | Pba (SRCI1043) *araBA*  ECARS11170, ECARS11175 | pWSK29 | 5’-CCTGCAGGGTGTAATTATCTTTTCATTCGC  5’-GGCTGCAGGCCAGTTACCTCGGTTCAAA |
| pJM068  pWSK*araF_Pba_::gfp+* | Pba (SRCI1043) *araF*  ECORS11165 | pWSK29 | 5’-GGCTGCAGGCCAGTTACCTCGGTTCAAA  5’-CCTGCAGGGTGTAATTATCTTTTCATTCGC |
| pTOF_ECs0066ko | ECs0066 | pTOF24 | ECs0066No_for: 5’-AAAAACTGCAGCGGTCGGTCGATAAAAAA  ECs0066Ni_rev: 5’-CGCTTCTTGCGGCCGCTTGGAACGGAAAGCTCGCACAGAATCA  ECs0066Ci-for: 5’-CCGTTCCAAGCGGCCGCAAGAGCGGCCAACTCACCGACTACT  ECs0066Co_rev: 5’-AAAAAGTCGACCATTTGATTGGCTGTGGTT |
| n/a | ECs0066 | qRT-PCR | araA-F.1: 5’-CGGTCACTGGCAGGATAAAC  araA-R.1: 5’-GACGGAGAAACCGAACTTGA |
| n/a | ECs0065 | qRT-PCR | araD-F.2: 5’-GCAAACGCTGCTGGATAAAC  araD-R.2: 5’-GCCTGGTTTCATTTGATTGG |

**Supplementary Table 2	Expression levels of arabinose reporters in xylose compared to arabinose**

|  | **Arabinose** | **Xylose** | **Xyl/Ara (%)** | **Glycerol** | **Gly/Ara (%)** |
| --- | --- | --- | --- | --- | --- |
| **pJM058** | 2066 (± 62) | 33 (± 6) | 1.60 | 4 (±4) | 0.20 |
| **pJM066** | 1574 (±26) | 528 (± 3) | 33.53 | 151 (±14) | 9.59 |
| **pJM064** | 14998 (±230) | 2994 (±71) | 19.96 | 1239 (±70) | 8.26 |
| **pJM065** | 31693 (±409) | 13584 (±640) | 42.86 | 8717 (±279) | 27.50 |

GFP was measured in response to 10 mM L-arabinose, 10 mM D-xylose or no added sugar (i.e. glycerol) at maximal expression times (four or six h), for pJM058 or pJM066 transformed in STEC (Sakai) or pJM064 / pJM065 (multi-copy plasmids) in Pba (1043), at 27 °C. The proportional of expression relative to arabinose is shown for xylose and glycerol as a percentage.

### Supplementary Table 3 Antibody screening of a panel of plant species, cultivars and tissues

|  |  |  |  | **Averaged absorbance for antibodies (OD 405nm)** | | | | | | **Standard deviation** | | | | | |
| --- | --- | --- | --- | --- | --- | --- | --- | --- | --- | --- | --- | --- | --- | --- | --- |
| **Species name** | **Common name** | **Cultivar** | **Tissue** | **LM6** | **LM13** | **LM5** | **LM16** | **LM1** | **JIM13** | **LM6** | **LM13** | **LM5** | **LM16** | **LM1** | **JIM13** |
| *Hordeum vulgare* | barley | Optic | leaf | 0.172 | 0.015 | 0.012 | 0.019 | 0.032 | 0.050 | 0.002 | 0.009 | 0.003 | 0.003 | 0.003 | 0.005 |
| *Ocimum basilicum* | basil | Gecofure | leaf | 0.080 | 0.003 | 0.039 | 0.009 | 0.002 | 0.365 | 0.002 | 0.001 | 0.010 | 0.005 | 0.001 | 0.008 |
| *Lactuca sativa* | lettuce | All Year Round | leaf | 0.053 | 0.038 | 0.128 | 0.015 | 0.006 | 0.048 | 0.008 | 0.012 | 0.014 | 0.001 | 0.002 | 0.016 |
| *Lactuca sativa* | lettuce | Butterhead | leaf | 0.017 | 0.012 | 0.127 | 0.008 | 0.006 | 0.050 | 0.003 | 0.004 | 0.005 | 0.004 | 0.004 | 0.003 |
| *Lactuca sativa* | lettuce | Little Gem | leaf | 0.021 | 0.018 | 0.247 | 0.005 | 0.003 | 0.035 | 0.001 | 0.009 | 0.021 | 0.004 | 0.003 | 0.002 |
| *Lactuca sativa* | lettuce | Rosetta | leaf | 0.009 | 0.018 | 0.105 | 0.009 | 0.003 | 0.050 | 0.002 | 0.006 | 0.043 | 0.003 | 0.001 | 0.002 |
| *Lactuca serriola* | wild lettuce | Serriola | leaf | 0.027 | 0.029 | 0.280 | 0.011 | 0.002 | 0.047 | 0.002 | 0.002 | 0.018 | 0.001 | 0.001 | 0.007 |
| *Raphanus sativus* | radish | Celesta | leaf | 0.087 | 0.054 | 0.048 | 0.009 | 0.024 | 0.012 | 0.015 | 0.022 | 0.019 | 0.005 | 0.002 | 0.002 |
| *Raphanus sativus* | radish | Expo | leaf | 0.119 | 0.046 | 0.014 | 0.012 | 0.029 | 0.028 | 0.013 | 0.006 | 0.001 | 0.007 | 0.002 | 0.001 |
| *Raphanus sativus* | radish | Sparkler | leaf | 0.111 | 0.130 | 0.028 | 0.015 | 0.047 | 0.020 | 0.017 | 0.026 | 0.000 | 0.007 | 0.001 | 0.011 |
| *Spinacia oleracea* | spinach | Amazon | leaf | 0.285 | 0.310 | 0.309 | 0.013 | 0.011 | 0.097 | 0.054 | 0.041 | 0.022 | 0.003 | 0.007 | 0.034 |
| *Spinacia oleracea* | spinach | Perpetual | leaf | 0.582 | 0.176 | 0.322 | 0.167 | 0.133 | 0.530 | 0.031 | 0.049 | 0.007 | 0.014 | 0.015 | 0.067 |
| *Spinacia oleracea* | spinach | Viking | leaf | 0.206 | 0.073 | 0.050 | 0.018 | 0.017 | 0.328 | 0.019 | 0.003 | 0.008 | 0.003 | 0.001 | 0.068 |
| *Solanum lycopersicum* | tomato | Alisa Craig | leaf | 0.044 | 0.018 | 0.052 | 0.010 | 0.016 | 0.437 | 0.002 | 0.003 | 0.009 | 0.004 | 0.001 | 0.079 |
| *Solanum lycopersicum* | tomato | Moneymaker | leaf | 0.048 | 0.015 | 0.042 | 0.009 | 0.012 | 0.496 | 0.013 | 0.004 | 0.000 | 0.002 | 0.003 | 0.073 |
| *Hordeum vulgare* | barley | Optic | root | 0.250 | 0.079 | 0.045 | 0.098 | 0.001 | 0.061 | 0.058 | 0.060 | 0.040 | 0.090 | 0.001 | 0.016 |
| *Ocimum basilicum* | basil | Gecofure | root | 0.252 | 0.037 | 0.030 | 0.019 | 0.004 | 0.220 | 0.020 | 0.000 | 0.005 | 0.013 | 0.001 | 0.020 |
| *Lactuca sativa* | lettuce | All Year Round | root | 0.295 | 0.158 | 0.500 | 0.014 | 0.018 | 0.198 | 0.016 | 0.070 | 0.033 | 0.006 | 0.016 | 0.060 |
| *Lactuca sativa* | lettuce | Butterhead | root | 0.069 | 0.016 | 0.128 | 0.009 | 0.013 | 0.021 | 0.006 | 0.008 | 0.001 | 0.003 | 0.006 | 0.010 |
| *Lactuca sativa* | lettuce | Little Gem | root | 0.092 | 0.017 | 0.144 | 0.018 | 0.002 | 0.038 | 0.008 | 0.008 | 0.004 | 0.005 | 0.001 | 0.014 |
| *Lactuca sativa* | lettuce | Rosetta | root | 0.101 | 0.009 | 0.120 | 0.005 | 0.003 | 0.026 | 0.023 | 0.005 | 0.023 | 0.000 | 0.003 | 0.005 |
| *Lactuca serriola* | wild lettuce | Serriola | root | 0.109 | 0.104 | 0.201 | 0.072 | 0.013 | 0.126 | 0.026 | 0.029 | 0.048 | 0.008 | 0.005 | 0.107 |
| *Raphanus sativus* | radish | Celesta | root | 0.115 | 0.020 | 0.012 | 0.015 | 0.036 | 0.039 | 0.003 | 0.008 | 0.001 | 0.004 | 0.004 | 0.004 |
| *Raphanus sativus* | radish | Expo | root | 0.487 | 0.017 | 0.007 | 0.022 | 0.016 | 0.122 | 0.007 | 0.002 | 0.007 | 0.008 | 0.001 | 0.005 |
| *Raphanus sativus* | radish | Sparkler | root | 0.257 | 0.023 | 0.070 | 0.036 | 0.058 | 0.095 | 0.023 | 0.012 | 0.033 | 0.018 | 0.046 | 0.048 |
| *Spinacia oleracea* | spinach | Amazon | root | 0.161 | 0.026 | 0.707 | 0.006 | 0.012 | 0.010 | 0.023 | 0.023 | 0.071 | 0.006 | 0.010 | 0.005 |
| *Spinacia oleracea* | spinach | Perpetual | root | 0.326 | 0.062 | 0.102 | 0.032 | 0.032 | 0.434 | 0.014 | 0.003 | 0.000 | 0.011 | 0.002 | 0.002 |
| *Spinacia oleracea* | spinach | Viking | root | 0.154 | 0.033 | 0.060 | 0.041 | 0.029 | 0.369 | 0.014 | 0.013 | 0.022 | 0.013 | 0.013 | 0.104 |
| *Solanum lycopersicum* | tomato | Alisa Craig | root | 0.306 | 0.221 | 0.256 | 0.142 | 0.120 | 0.541 | 0.018 | 0.021 | 0.080 | 0.008 | 0.008 | 0.104 |
| *Solanum lycopersicum* | tomato | Moneymaker | root | 0.429 | 0.163 | 0.222 | 0.214 | 0.173 | 0.478 | 0.020 | 0.056 | 0.036 | 0.026 | 0.008 | 0.060 |
| *Brassica oleracea var. italica* | broccoli* | Marathon | microleaf | 0.354 | 0.014 | 0.008 | 0.038 | 0.009 | 0.421 | 0.075 | 0.000 | 0.001 | 0.003 | 0.004 | 0.086 |
| *Medicago sativa* | alfalfa | not known | sprout | 0.281 | 0.245 | 0.107 | 0.122 | 0.109 | 0.155 | 0.021 | 0.038 | 0.003 | 0.023 | 0.031 | 0.045 |
| *Trigonella foenum-graecum* | fenugreek | not known | sprout | 0.313 | 0.258 | 0.060 | 0.061 | 0.048 | 0.057 | 0.005 | 0.012 | 0.005 | 0.001 | 0.003 | 0.005 |

Pectin-enriched fractions (CDTA-treated, as per [1]) of horticultural and arable crop leaves and roots (barley, basil, lettuce, wild lettuce, radish, spinach, tomato), and sprouted seeds (alfalfa, fenugreek, prepared as per [2]) screened with selected antibodies (LM6, anti-(1→5)-α-L-arabinan (1 residue) [3]; LM13, anti-(1→5)-α-L-arabinan (3+ residues) [4]; LM5, anti-(1→4)-β-D-galactan [5]; LM16, Rhamnogalacturonan I epitope (uncharacterised) [4]; LM1, Extensin (histidine-rich glycoprotein) [6]; JIM13, β-D-GlcA-(1,3)-α-D-GalA-(1,2)-α-L-Rha (arabinogalactan protein) [7]). ELISA data is expressed as averaged absorbance (405 nm) in a heat-map format with standard deviations from replicated samples. ELISA data colour scale: high (red) to low (white). * broccoli microgreen data added from Fig. 5 for comparison, probed with the same antibodies except LM2 (1 → 6)-β-D galactan chain with terminally attached GlcA (arabinogalactan protein) [6, 7] was used in place of JIM13.
